# UFMylation promotes orthoflavivirus infectious particle production

**DOI:** 10.1101/2025.01.09.632082

**Authors:** Hannah M. Schmidt, Grace C. Sorensen, Matthew R. Lanahan, Jenna Grabowski, Moonhee Park, Stacy M. Horner

## Abstract

Post-translational modifications play crucial roles in viral infections, yet many potential modifications remain unexplored in orthoflavivirus biology. Here we demonstrate that the UFMylation system, a post-translational modification system that catalyzes the transfer of UFM1 onto proteins, promotes infection by multiple orthoflaviviruses including dengue virus, Zika virus, West Nile virus, and yellow fever virus. We found that depletion of the UFMylation E3 ligase complex proteins UFL1 and UFBP1, as well as other UFMylation machinery components (UBA5, UFC1, and UFM1), significantly reduces infectious virion production for orthoflaviviruses but not the hepacivirus, hepatitis C. Mechanistically, UFMylation does not regulate viral RNA translation or RNA replication but instead affects a later stage of the viral lifecycle. We identified novel interactions between UFL1, and several viral proteins involved in orthoflavivirus virion assembly, including NS2A, NS2B-NS3, and Capsid. These findings establish UFMylation as a previously unrecognized post-translational modification system that promotes orthoflavivirus infection, likely through modulation of viral assembly. This work expands our understanding of the post-translational modifications that control orthoflavivirus infection and identifies new potential therapeutic targets.

**Importance:** Orthoflaviviruses depend on host-mediated post-translational modifications to successfully complete their lifecycle, yet many of these critical interactions remain undefined. Here, we describe a role for a post-translational modification pathway, UFMylation, in promoting infectious particle production of ZIKV and DENV. We show that UFMylation regulates these viruses at a lifecycle stage after initial RNA translation and RNA replication. Additionally, we find that regulation of infection by UFMylation extends to other orthoflaviviruses, including West Nile virus and yellow fever virus, but not to the broader Flaviviridae family. Finally, we demonstrate that UFMylation machinery directly interacts with specific DENV and ZIKV proteins during infection. These studies reveal a previously unrecognized role for UFMylation in regulating orthoflavivirus infection.

## Introduction

Orthoflaviviruses are a genus of positive-sense RNA viruses (1) that represent a significant human health burden. These viruses, which include dengue virus (DENV), West Nile Virus (WNV), yellow fever virus (YFV), and Zika virus (ZIKV), are transmitted by arthropods in tropical regions, placing billions of people at risk of contracting an orthoflavivirus infection annually (2). There are currently a lack of therapies and broadly effective vaccines against these viruses, highlighting the need for a better understanding of the molecular processes occurring during viral infection. Orthoflaviviruses have a compact but efficient genome organization and lifecycle that enables successful viral infection. Within infected cells, the ∼11 kilobase positive-sense RNA genome is translated as a single polyprotein that is cleaved by viral and host proteases into ten individual proteins (3). These viral proteins include three structural proteins (C, prM/M, and E), which form the virion, and seven non-structural proteins (NS1, NS2A, NS2B, NS3, NS4A, NS4B, and NS5) that mediate viral replication and coordinate additional functions that contribute to infection, such as facilitating evasion of the innate immune system (4). Following translation, the viral proteins induce ER invaginations to compartmentalize viral RNA replication (3). Then, the viral genomic RNA is transported from the ER invaginations to associate with the viral Capsid (C) protein, forming a nucleocapsid (5). The viral nucleocapsid buds through the ER to form an immature virion which undergoes additional maturation before being secreted from the cell (3). Due to their limited genome size, orthoflaviviruses rely on host factors to dynamically regulate the roles of viral proteins in distinct viral lifecycle stages (6–8). While many roles for host proteins in orthoflavivirus infection have been characterized, the full scope of host factors regulating orthoflavivirus infection is unknown, including roles for many enzymes catalyzing post-translational modifications.

Viral infection can be regulated by reversible post-translational modifications (9), which modify viral or host proteins to alter their stability, subcellular localization, and function. While roles for some post-translational modifications of orthoflaviviral proteins, such as acetylation, phosphorylation, ubiquitination, and glycosylation, have been described (10–13), there are many post-translational modifications, including novel ubiquitin-like modifications, that have not been fully explored during orthoflavivirus infection. One such ubiquitin-like modification is UFM1. UFM1 is an 85-amino acid ubiquitin-like peptide conjugated onto lysine residues through an enzymatic pathway involving the E1 activase, UBA5, the E2 conjugase, UFC1, and the E3 ligase complex, UFL1-UFBP1, while UFSP2 mediates its removal (14–19). The addition of UFM1 to proteins, which is referred to as UFMylation, can regulate protein function by altering protein-protein interactions (20–22). UFMylation regulates several host processes essential to viral infection and can modulate infection by a number of diverse viruses, including the gamma-herpesvirus Epstein-Barr virus (EBV) (23) and the picornavirus hepatitis A virus (HAV) (24), ultimately limiting inflammation or promoting viral translation during infection, respectively. UFL1 has also been described to regulate pathways that could impact orthoflavivirus infection, such as promoting antiviral RIG-I signaling (25) and resolving ER stress responses (20, 26, 27), which are known to accumulate during orthoflavivirus replication (28). However, a specific function for UFMylation during orthoflavivirus infection has not been described.

Here, we demonstrate that UFL1, the UFMylation machinery, and the process of UFM1 conjugation promote DENV and ZIKV infectious virion production through a mechanism independent of viral RNA translation or replication. Additionally, we find that UFL1 promotes infection of several orthoflaviviruses, including DENV, ZIKV, WNV, and YFV, but it does not regulate infection by the hepacivirus, hepatitis C virus (HCV). Mechanistically, we find that UFL1 can interact with the viral Capsid, NS2A, and NS2B/NS3 proteins during orthoflavivirus infection, suggesting that UFL1 interactions with these viral proteins may regulate their function through direct modification or altered protein-protein interactions. These findings establish UFM1 and the process of UFMylation as post-translational regulators of orthoflavivirus infection.

## Results

### The UFMylation E3 ligase complex proteins promote orthoflavivirus infection

To determine if the UFMylation E3 ligase complex protein UFL1 regulates orthoflavivirus infection, we examined the production of infectious virions during DENV or ZIKV infection following siRNA-mediated depletion of UFL1 in human hepatoma Huh7 cells. Huh7 cells are an appropriate model cell line for these viruses because they can support high levels of orthoflavivirus infection. In addition, orthoflavivirus infection induces disease pathologies associated with viral infection in the liver (29). We found that compared to cells treated with control non-targeting siRNA, depletion of UFL1 decreased the levels of infectious DENV in the supernatant at 48 hours post infection, as measured by focus-forming assay (Figure 1A, left). Similarly, depletion of UFL1 reduced the levels of infectious ZIKV (Figure 1A, right). As recent studies have shown that the E3 ligase of UFMylation is a complex consisting of both UFL1 and UFBP1 (17, 30) we next examined the role of UFBP1 in DENV and ZIKV infection. We found that depletion of UFBP1 in Huh7 cells also resulted in decreased levels of infectious virions from both DENV and ZIKV infections, indicating that the UFMylation E3 ligase complex promotes orthoflavivirus infection (Figure 1B). Importantly, depletion of UFL1 or UFBP1 did not affect the viability of Huh7 cells (Figure 1C). However, the loss of expression of either UFL1 or UFBP1 in Huh7 cells did result in reduced expression of its cognate cofactor (Figure 1D), as seen previously by others (17, 31). We observed similar results in A549 cells, a lung carcinoma line susceptible to orthoflavivirus infection, where depletion of either UFL1 or UFBP1 decreased the production of infectious ZIKV virions and resulted in reduced expression of both proteins (Figure 1E). These results confirm that depletion of either UFL1 or UFBP1 results in loss of the overall UFMylation E3 ligase complex. In addition, these data reveal that depletion of the UFMylation E3 ligase complex results in reduced viral particle production during DENV and ZIKV infection. Moving forward, we solely targeted UFL1 expression to manipulate expression of the UFMylation E3 ligase complex.

**Figure 1.**
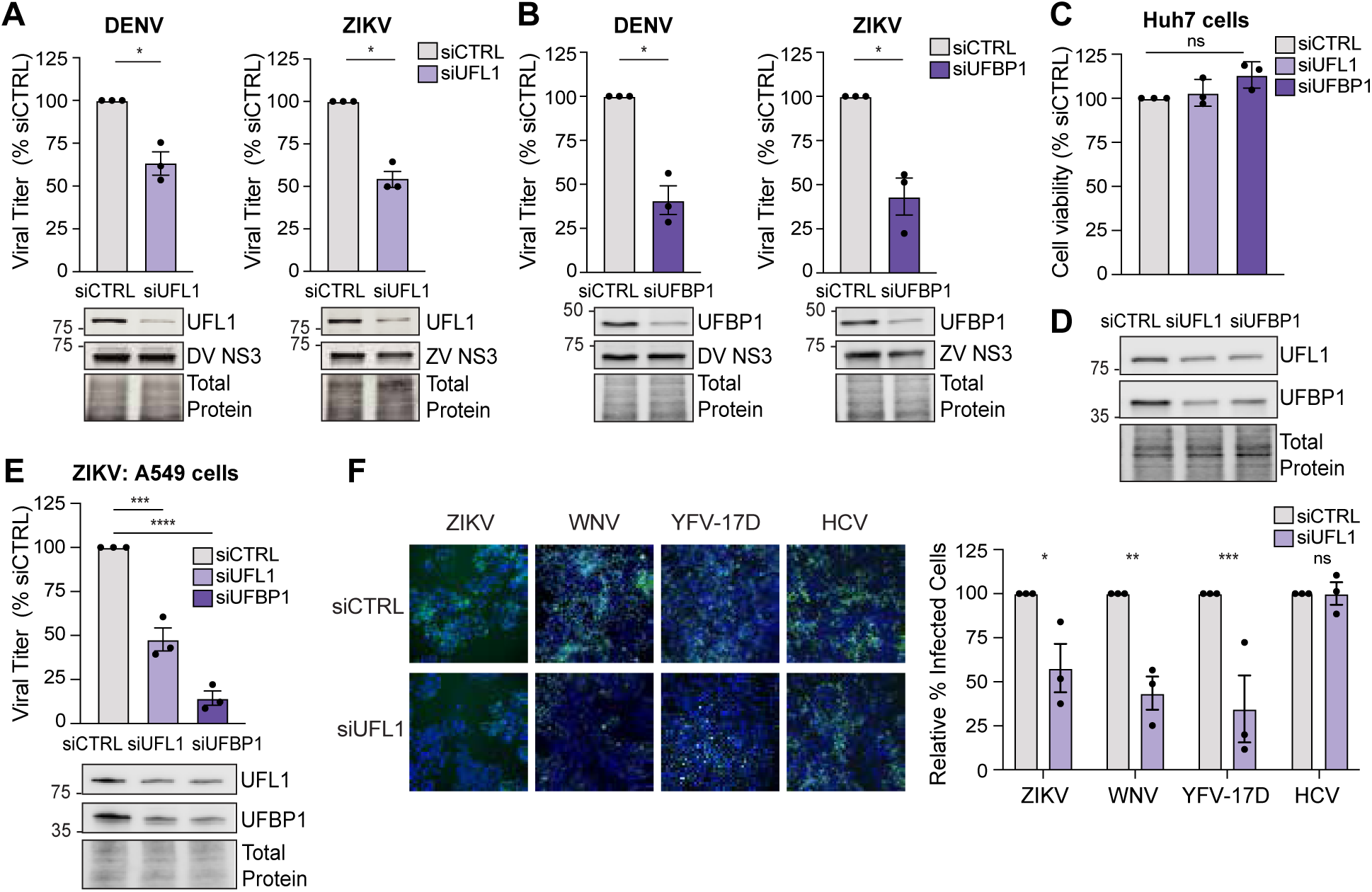
The UFMylation E3 ligase complex proteins promote mosquito-borne orthoflavivirus infection. (**A** and **B**) Focus-forming assay of supernatants from Huh7 cells infected with DENV^NGC^ or ZIKV^PRVABC59^ (48 h, MOI 0.1) after siRNA depletion of the indicated transcripts or non-targeting control (CTRL), shown as % of siCTRL. (**C**) Cell viability measured after siRNA depletion of the indicated transcripts at 72 hours post-transfection, relative to that of siCTRL, as measured by Cell-Titer GLO assay. **(D**) Immunoblot analysis of protein expression from Huh7 cells treated with the indicated siRNAs for 72 hours. (**E**) Focus-forming assay of supernatants harvested from A549 cells infected with ZIKV^PRVABC59^ (48 h, MOI 0.1) after siRNA depletion of the indicated transcripts. (**F**) Immunofluorescence micrographs of Huh7 cells treated with indicated siRNA and then infected with the following viruses for 48 hours (ZIKV^PRVABC59^, MOI 0.1; YFV^17D^, MOI 0.01; WNV^NY2000^, MOI 0.01, or HCV^JFH1^, MOI 1), as measured by immunostaining of viral antigen (E for ZIKV, YFV^17D^, and WNV, NS5A for HCV; green). Nuclei were stained with Hoechst (blue). Right: Quantification of the percentage of virus-infected Huh7 cells, shown relative to siCTRL. >5000 cells counted for each condition. For all panels, n=3 biologically independent experiments, with bars indicating mean and error bars showing standard error of the mean. *p<0.05, ***p<0.001, ****p<0.0001, or ns, not significant as determined by paired t-test (A and B), one-way ANOVA with Dunnett’s multiple comparisons test (C and E), or two-way ANOVA followed by Šidák’s multiple comparisons test (F).

As the UFMylation E3 ligase complex was required to promote infection by both DENV and ZIKV, we next tested if it also regulated infection by other viruses in the *Flaviviridae* family. We depleted UFL1 by siRNA in Huh7 cells and measured the percent of virus-infected cells during WNV, YFV^17D^, or HCV infection, using ZIKV as our control virus, as we found it was regulated by UFL1. We found that UFL1 depletion resulted in a ∼50% decrease in the percentage of cells infected by the orthoflaviruses ZIKV, WNV, and YFV-17D (Figure 1F). However, UFL1 depletion had no effect on percentage of cells infected by the hepacivirus HCV (Figure 1F).Taken together, these data indicate that the UFMylation E3 ligase complex promotes infection of several orthoflaviviruses, but not all viruses in the *Flaviviridae* family.

### The UFMylation E3 ligase complex does not regulate orthoflavivirus translation or RNA replication

Having found that UFL1 and UFBP1 regulate orthoflavivirus infection, we next wanted to map the stage of the orthoflavivirus life cycle regulated by the UFMylation E3 ligase complex, using ZIKV and DENV as representative orthoflaviviruses. As others have shown that UFL1 and UFBP1 promote translation of the genome of the positive-strand RNA virus hepatitis A virus (24), we tested if UFL1 is required for ZIKV RNA translation. To do this, we first established timepoints corresponding to initial RNA translation and replication during infection with an infectious ZIKV reporter virus that encodes *Gaussia* luciferase (ZIKV-GLuc) (32), measuring *Gaussia* luciferase activity over time. We detected *Gaussia* luciferase activity as early as 3 hours post-infection, and this activity increased over the time course from 3 to 12 hours, indicative of increased *Gaussia* luciferase expression (Figure 2A). Importantly, cycloheximide treatment, which inhibits translation, resulted in decreased ZIKV-GLuc levels as early as 3 hours post-treatment (Figure 2A), while MK0608 treatment, which inhibits the viral RdRp (33), resulted in decreased ZIKV-GLuc levels at time points later than 9 hours post-treatment (Figure 2A). These results demonstrate that the signal observed from ZIKV-GLuc at 3 and 6 hours is the product of viral translation in itself, and that following 9 hours, viral RNA replication also contributes to the increasing levels of ZIKV-GLuc. Having established this system to measure the translation and replication of ZIKV, we next determined if UFL1 depletion alters these viral lifecycle steps. To do this, we depleted UFL1 by siRNA in Huh7 cells, infected with ZIKV-GLuc, and measured GLuc expression over time. In the 3-12 hours post-infection, which we established measures RNA translation and replication of ZIKV, depletion of UFL1 had no effect on the relative ZIKV-GLuc levels (Figure 2B). However, depletion of UFL1 did reduce the levels of ZIKV-GLuc in the 24-72 hours after infection (Figure 2C-2D). This suggests that UFL1 does not regulate viral RNA translation or RNA replication and instead acts on a later viral lifecycle stage.

**Figure 2.**
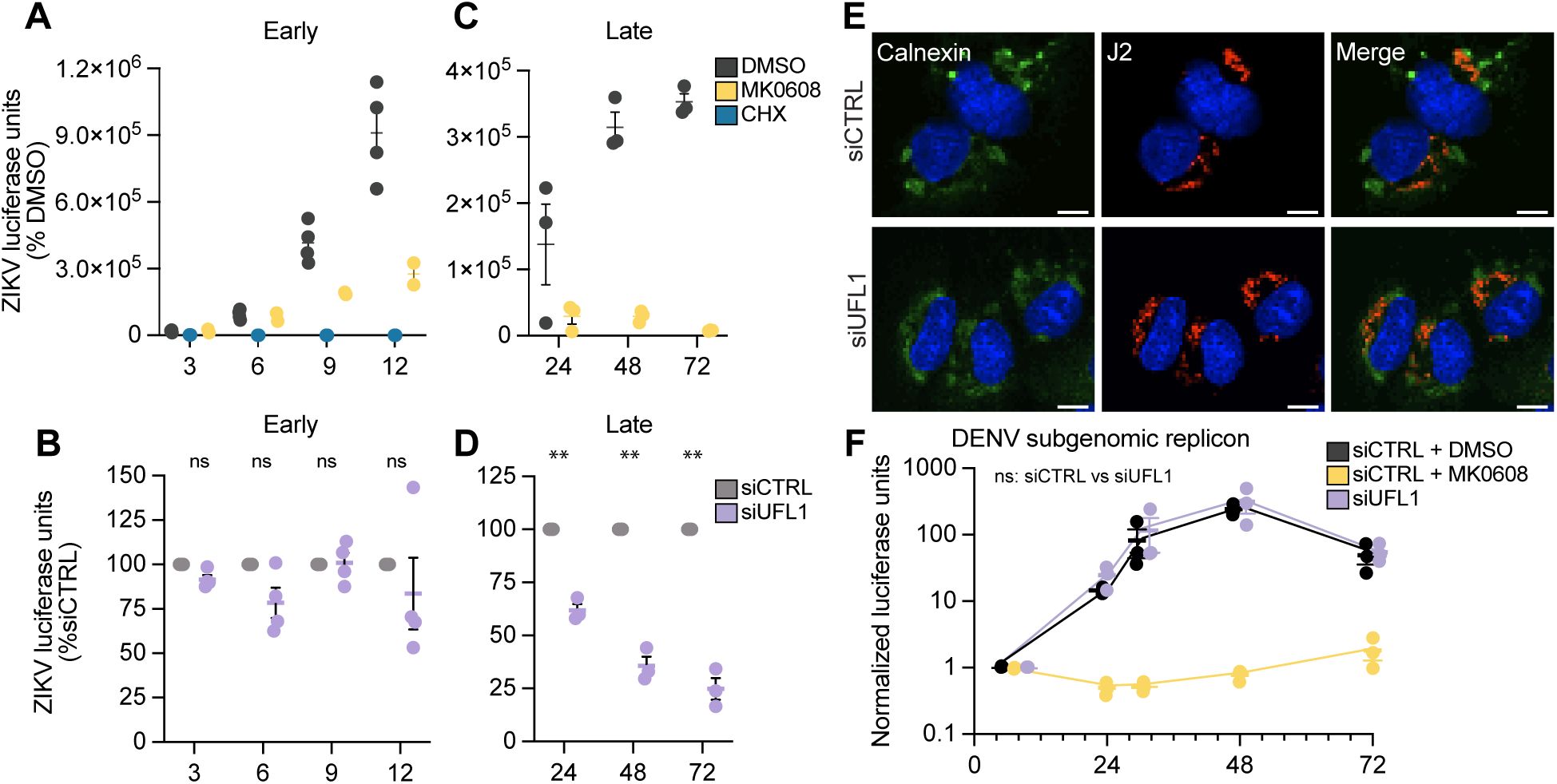
The UFMylation E3 ligase complex does not regulate orthoflavivirus translation or RNA replication. (**A)** Luciferase activity of *Gaussia* luciferase-encoding ZIKV^MR766^ (ZIKV-GLuc, MOI 0.1) from infected Huh7 cells treated with DMSO, MK0608, or cycloheximide during infection and harvested at the indicated time points. (**B**) Normalized expression of ZIKV-GLuc from infected Huh7 cells treated with non-targeting control (CTRL) or UFL1 siRNA harvested at the indicated timepoints. **(C)** Luciferase activity of *Gaussia* luciferase-encoding ZIKV^MR766^ (ZIKV-GLuc, MOI 0.1) from supernatant of infected Huh7 cells treated with DMSO or MK0608 harvested at 24, 48, or 72 hpi. (**D)** Normalized luciferase activity of ZIKV-GLuc from supernatant of infected Huh7 cells treated with CTRL or UFL1 siRNA and harvested at 24, 48, or 72 hpi.(**E**) Immunofluorescence micrographs of Huh7 cells treated with the indicated siRNA and then infected with ZIKV^PRVABC59^ (36 h, MOI 1) that were immunostained with anti-calnexin (green) and anti-J2 (red) for dsRNA, with the nuclei stained with Hoechst (blue). Scale bar, 10 μm. (**F**) Normalized luciferase expression of lysates from expression of Huh7 cells transfected with the indicated siRNA and electroporated with a DENV^16681^ subgenomic RNA replicon expressing *Renilla* luciferase harvested at the indicated timepoints. Treatment with MK0608 was as in (**A**). For all panels, n=3 biologically independent experiments, with bars indicating mean and error bars showing standard error of the mean. **p<0.01, or ns, not significant, determined by two-way ANOVA with Dunnett’s multiple comparisons test (B, D, and F)

After the initial rounds of orthoflavivirus RNA translation, the viral proteins induce ER invaginations that compartmentalize viral RNA replication (34). As the UFMylation E3 ligase complex can regulate ER morphology (31), we next tested if UFL1 regulates the general morphology of the ER in ZIKV-infected Huh7 cells by examining the gross morphology of the characteristic viral dsRNA-containing ER membranes that accumulate in the perinuclear region, as seen by others (35, 36). Using immunofluorescence morphology with staining for the ER (Calnexin) and dsRNA (J2), we found that depletion of UFL1 did not appear to broadly alter this morphology (Figure 2E). To directly test if UFL1 regulates orthoflavivirus replication, we measured the replication of a subgenomic RNA replicon of DENV encoding a *Renilla* Luciferase gene (DENV-RLuc-SGR) (8). This subgenomic RNA replicon lacks the viral structural genes but contains the non-structural genes sufficient for RNA replication, such that when *in vitro* transcribed RNA is transfected into cells, the viral RNA can replicate but cannot produce infectious virions. Following transfection of *in vitro* transcribed DENV-RLuc-SGR RNA into Huh7 cells, we found that UFL1 depletion did not alter RLuc activity levels over a time course, while the RdRp inhibitor MK0608 did prevent RLuc expression, as expected, because it inhibits RNA replication (Figure 2F). In summary, these data reveal that UFL1, and thus the UFMylation E3 ligase complex, promotes orthoflavivirus infection at a viral lifecycle stage following RNA replication.

### The UFMylation machinery promotes orthoflavivirus infection

Having found that the UFMylation E3 ligase complex promotes orthoflavivirus infection, we next wanted to determine if the other proteins that regulate UFMylation beyond the E3 ligase complex promote infection by DENV and ZIKV. To test this, we depleted the E1 activase UBA5, the E2 conjugase UFC1, or the ubiquitin-like modifier UFM1 in Huh7 cells using siRNA and then infected these cells with ZIKV and DENV. Importantly, we validated knockdown of the proteins by immunoblotting and confirmed that transient depletion of the UFMylation machinery did not affect cell viability of Huh7 cells compared to siCTRL as measured by Cell-Titer GLO assay (Figure 3A and 3B). Depletion of each of the UFMylation machinery proteins reduced infectious virion production of ZIKV between ∼35-65% compared to a non-targeting control (Figure 3C). Similarly, depletion of the UFMylation machinery proteins reduced the infectious virion production of DENV (Figure 3D). Importantly, we also validated our earlier results showing that depletion of UFL1 by siRNA resulted in reduced infectious virion production in either ZIKV or DENV (Figure 3C and 3D). Since the complement of proteins involved in the process of UFM1 conjugation all positively regulate ZIKV and DENV infectious virion production, we next wanted to test if UFM1 conjugation itself is required to promote ZIKV and DENV infection. To do this, we transduced Huh7-UFM1 KO cells that we generated by CRISPR/Cas9 with lentiviruses expressing Flag-tagged UFM1^WT^ or UFM1^ΔC3^, in which the deletion of the last three residues of UFM1 prevents its conjugation to the lysine residues of target proteins (15). Importantly, we confirmed that UFM1^ΔC3^ limits UFM1 conjugation, while UFM1^WT^ maintains UFM1 conjugation, by measuring the formation of UFM1-conjugates by immunoblotting in Huh7-UFM1 KO cells complemented with Flag-UFM1^ΔC3^ (Figure 3E and 3F). When we infected with ZIKV, we found that Huh7-UFM1 KO cells complemented with FLAG-UFM1^ΔC3^ produced roughly half as many infectious virions as those cells complemented with FLAG-UFM1^WT^ (Figure 3E). While the mean production of infectious DENV virions was lower in the Flag-UFM1^ΔC3^ cells, this decrease was not statistically significant (Figure 3F). Taken together, these data indicate that the UFMylation machinery proteins and UFM1 conjugation itself promote orthoflavivirus infection.

**Figure 3.**
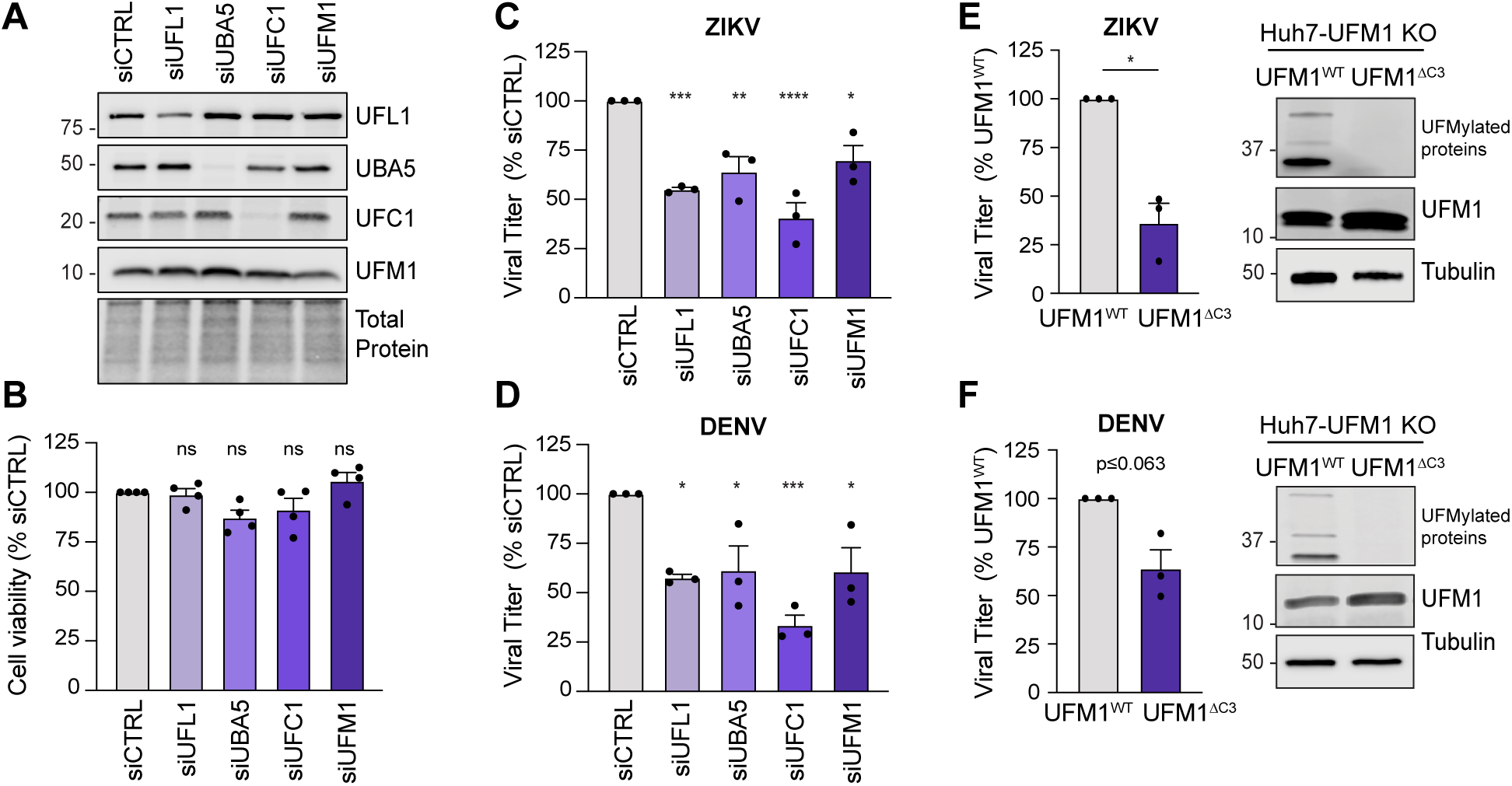
The UFMylation machinery promotes orthoflavivirus infection. (**A**) Immunoblot analysis of Huh7 cells after siRNA depletion of the indicated transcripts or non-targeting control (CTRL). (**B)** Cell viability measured after siRNA depletion of the indicated transcripts at 72 hours post-transfection, as measured by Cell-Titer GLO assay, relative to the viability of siCTRL. (**C**-**D**) Focus-forming assay of supernatants harvested from Huh7 cells infected with DENV^NGC^ or ZIKV^PRVABC59^ (48 h, MOI 0.1) after siRNA depletion of the indicated transcripts, shown as % of siCTRL. (**E-F**) Focus-forming assay of supernatants harvested from Huh7-UFM1 KO cells transduced with Flag-UFM1^WT^ or Flag-UFM1^ΔC3^ and infected with either DENV^NGC^ (72 h, MOI 0.1) or ZIKV ^PRVABC59^ (48 h, MOI 0.1), shown as % of Flag-UFM1^WT^. Immunoblots indicate UFM1-conjugated proteins as those that are higher molecular weight from unconjugated UFM1 but are detected with the anti-UFM1 antibody. For all panels, n=3 biologically independent experiments, with bars indicating mean and error bars showing standard error of the mean. *p<0.05, ** p<0.001, ***p<0.001, ****p<0.0001, or ns, not significant, determined by one-way ANOVA with Dunnett’s multiple comparisons test (B, C, and D) or paired t-test (E and F).

### UFL1 interacts with several DENV and ZIKV proteins

Our results so far have revealed that UFL1 promotes DENV and ZIKV infection at a viral lifecycle step that occurs following the initial viral RNA translation and RNA replication to promote infectious particle production. Several host factors are known to promote infectious particle production of orthoflaviviruses by interacting with viral proteins (5, 37–40). To uncover the mechanisms of how the UFMylation E3 complex regulates orthoflaviviral infectious virion production, we tested if UFL1 interacts with any of the orthoflaviviral proteins (Figure 4A). We focused on UFL1 because, unlike UFBP1, it is not anchored to ER membranes (31), allowing us to more easily define protein-protein interactions between viral proteins and the UFMylation E3 complex in co-immunoprecipitation-based experiments. In an initial screen, we measured the interaction of UFL1 with a V5-tagged set of DENV proteins (41) by co-immunoprecipitation in Huh7 cells. For the experiments, we utilized different lysis conditions depending on the viral protein. We found that DENV Capsid, NS2A, and the NS2B-NS3 complex can interact with UFL1 in an over-expression setting (Figure 4B).

**Figure 4.**
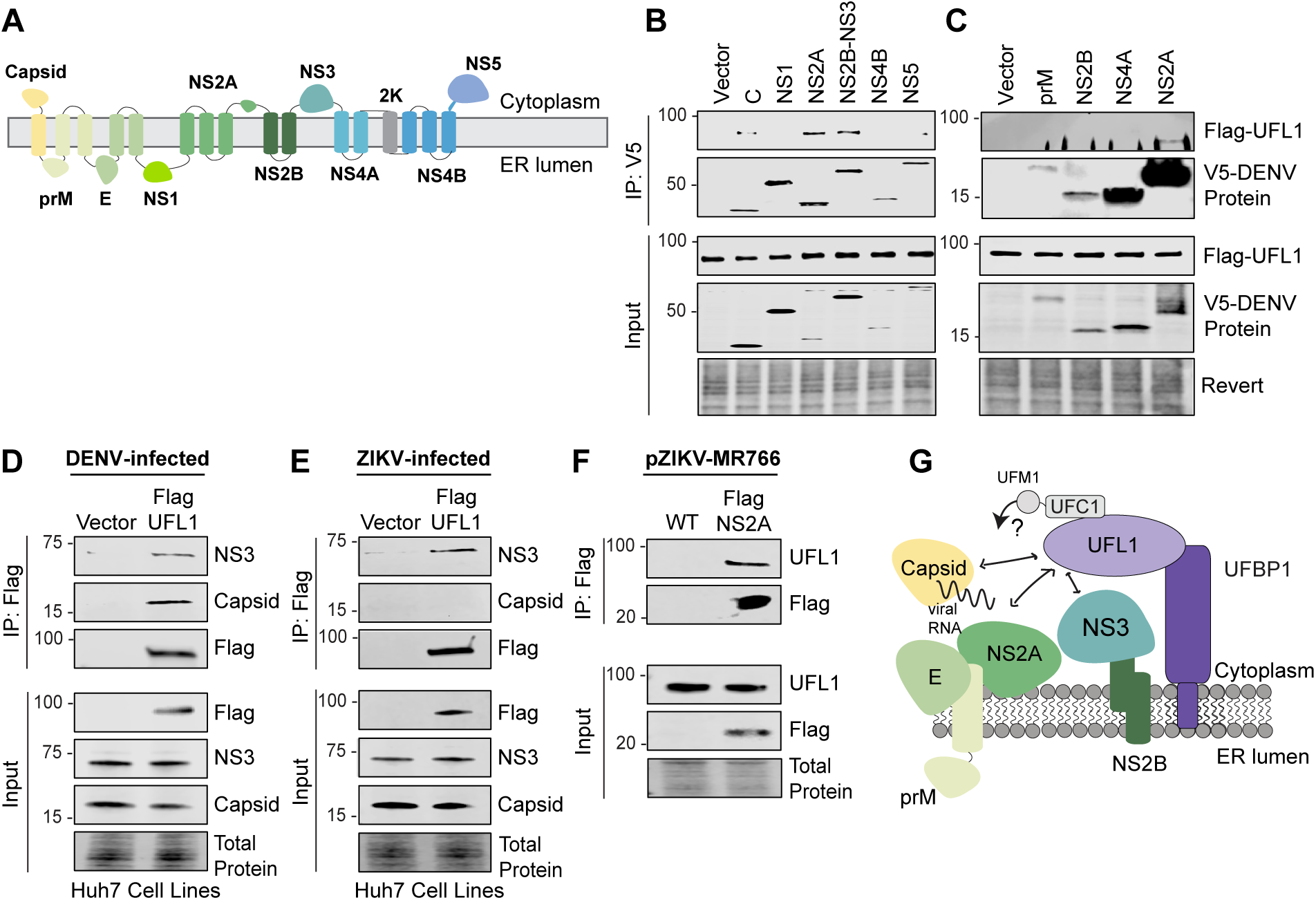
UFL1 interacts with several DENV and ZIKV proteins. (**A**) Schematic of DENV polyprotein, showing membrane topology of viral proteins. (**B-C**) Immunoblot analysis of anti-V5 immunoprecipitated extracts and inputs, lysed in NP40 buffer (**B**) or TX-100-RIPA buffer (**C**), from Huh7 cells stably expressing Flag-UFL1 transfected with plasmids expressing V5-tagged DENV^16681^ proteins. (**D-E**) Immunoblot analysis of anti-Flag immunoprecipitated extracts and inputs from DENV^NGC^-infected or ZIKV^PRVABC59^-infected (48 h, MOI 1) Huh7 cells stably expressing Flag-UFL1 or Vector. (**F**) Immunoblot analysis of anti-Flag immunoprecipitated extracts and inputs from Huh7 cells transfected with DNA plasmids encoding the plasmid-launched ZIKV^MR766-WT^ or pZIKV ^MR766-Flag-NS2A^ and harvested at 72 hpi. Representative immunoblots from n=3 biologically independent experiments are shown.

However, we did not detect interaction between UFL1 and prM, NS1, NS2B, NS4A, NS4B, or NS5 (Figure 4B and 4C). Of note, the interaction of NS2A with UFL1 was detected regardless of the lysis buffer. We did not screen for interaction with viral E protein as we found that this construct did not express and is unlikely to interact with the cytosolic UFL1, as it is localized to the lumen of the ER (3). Thus, over-expression-based co-immunoprecipitation assays suggest interaction between UFL1 and three DENV proteins: Capsid, NS2A, and NS2B-NS3.

We next measured these protein-protein interactions in the context of viral infection. We infected Huh7 cells stably expressing Flag-UFL1 or Flag-tag alone with DENV or ZIKV and immunoprecipitated Flag-UFL1. During DENV infection, we found that both NS3 and Capsid interact with UFL1 (Figure 4D). However, during ZIKV infection, we found that NS3, but not Capsid, interacts with UFL1 (Figure 4E). As no commercial antibodies are available for the orthoflaviviral NS2A protein, we sought to validate the interaction of UFL1 and NS2A during ZIKV infection using a ZIKV^Flag-NS2A^ expressing virus. This virus is similar to ones generated by others, where a protein tag is cloned into the junction between NS1 and NS2A (42). In this case a 3XFlag tag was inserted into the backbone of the plasmid-based rescue system for ZIKV MR766 (43). At 72 hours post-transfection of pZIKV^Flag-NS2A^ or pZIKV^WT^, which does not contain any epitope tag on NS2A and served as our negative control, we immunoprecipitated Flag-NS2A using an anti-Flag antibody and found that it co-immunoprecipitated with endogenous UFL1 (Figure 4F). Together, these data show that UFL1 interacts with specific DENV and ZIKV proteins during infection.

## Discussion

Orthoflavivirus infection is a tightly coordinated process regulated by multiple mechanisms, including the localization of viral proteins to replication complexes, ER membrane rearrangements, the stability of viral proteins, and post-translational modifications to both viral and host proteins (3, 6, 10). While post-translational modification of both viral and host proteins can regulate different aspects of the orthoflavivirus lifecycle (10–12, 44, 45), our understanding of the full complement of post-translational modifications that regulate infection by these viruses remains incomplete. Here, we have identified UFMylation as a post-translational modification system that positively regulates orthoflavivirus infection. Our data demonstrate that the UFMylation E3 ligase complex proteins, UFL1 and UFBP1, along with the broader UFMylation machinery, promote both DENV and ZIKV infectious particle production. In addition, we found that the UFMylation E3 ligase complex promotes infection by several different orthoflaviviruses, including DENV, ZIKV, WNV, and YFV. Mechanistically, we found that the UFMylation E3 ligase complex does not affect DENV and ZIKV RNA translation, genomic RNA replication, or the production of replication complexes, but instead regulates a late stage of the viral lifecycle that culminates in the production of infectious virions. Supporting this conclusion, we identified protein-protein interactions between UFL1 and orthoflaviviral proteins that can have roles in virion assembly, including NS2A and NS2B-NS3 for both DENV and ZIKV, and Capsid for DENV. Taken together, our results reveal a new role for the process of UFMylation in promoting infectious virion production during orthoflavivirus infection.

The process of UFMylation has been described to regulate infection by other viruses via diverse mechanisms. For example, during EBV infection, the viral protein BILF1 promotes the UFMylation of MAVS, which ultimately facilitates sorting of UFMylated MAVS into mitochondrial-derived vesicles for lysosomal degradation (23). During HAV infection, UFMylation of the ribosomal protein RPL26 promotes viral translation (24). The authors of this study speculated that the UFMylation of RPL26, which is located near the ribosome exit tunnel, may result in increased viral translation by resolving viral RNA structures that otherwise would limit viral translation (24). UFMylation has also been shown to regulate the antiviral innate immune response by facilitating RIG-I interaction with its known regulator 14-3-3ε to promote interferon induction (25). While these studies show that UFMylation regulates host protein functions during viral infection, our work differs in that it implicates UFL1 in modulating viral protein functions through interaction with NS2A, NS2B-NS3, and Capsid. This, combined with the result that the process of UFMylation regulates orthoflaviviral infection, raises the possibility that one or more of these viral proteins is UFMylated to promote infectious virion production. As UFMylation can regulate protein sorting or specific protein-RNA interactions in the viral infection systems described above (23, 24), it seems likely that during orthoflavivirus infection, UFMylation could regulate protein trafficking or protein-RNA interactions that coordinate viral RNA packaging into nascent virions.

Our understanding of the factors that regulate the production of infectious virions during orthoflavivirus infection is still incomplete. It is interesting that we found that UFL1 interacts with proteins from DENV and ZIKV that regulate aspects of viral assembly, specifically NS2A, NS2B-NS3, and Capsid proteins, suggesting that UFL1 and UFMylation may be regulating their function during assembly. We know that during orthoflavivirus assembly, NS2A appears to play a key role in bringing the viral RNA genome from the replication complex to the virion assembly site at the ER membrane (46–48). There, it also interacts with NS2B-NS3, which works with the host signal peptidase to cleave the Capsid-prM-E polyprotein and produce the individual viral Capsid, prM, and E proteins (5, 49). This allows NS2A to transfer the positive-strand genomic viral RNA to the Caspid protein, which oligomerizes to form the immature viral nucleocapsid, which then undergoes further maturation in the Golgi (5, 50) to produce a fully mature virion (51, 52). We hypothesize that UFMylation promotes one or more of the following steps of virion assembly (Figure 4G). These could be either (1) NS2A binding to viral RNA, (2) NS2A interactions with NS2B-NS3 for processing of the immature Capsid-prM-E polyprotein, or (3) Capsid RNA binding and nucleocapsid formation. Interestingly, as the interaction of UFL1 with Capsid protein occurs in DENV infection but not in ZIKV infection, this suggests that the mechanism by which UFMylation regulates orthoflavivirus virion assembly may be somewhat distinct between these two related viruses. Furthermore, while the interactions between UFL1 and several viral proteins suggest that the UFMylation machinery regulates viral infection through one of these proteins, we have not ruled out the possibility that UFMylation may also be regulating host processes that promote viral infection. Future work will be aimed at determining the UFMylation status of Capsid, NS2A, and NS2B-NS3, as well as characterizing which aspect of viral assembly may be modulated by UFMylation.

Post-translational modifications have emerged as direct modifiers of viral proteins, regulating multiple aspects of the orthoflaviviral life cycle. For example, K63-linked ubiquitination of ZIKV envelope promotes viral entry (12) and SUMOylation promotes DENV NS5 stability to facilitate replication (53). Our work adds the process of UFMylation to the complement of post-translational modification systems that regulate orthoflavivirus infection, alongside acetylation, glycosylation, phosphorylation, ubiquitination, and SUMOylation (10–13, 53), all of which may be avenues for anti-viral therapies. While targeting virus-host interactions remains a promising antiviral drug strategy, much remains to be learned about how the UFMylation machinery selects its targets to minimize the effects on host cell pathways. Altogether, our work here reveals that UFM1 is a novel post-translational regulator of orthoflavivirus infection, broadening our understanding of the host factors required to promote viral infection.

## Acknowledgements

We thank those colleagues who generously provided reagents, as indicated in the Methods; the Duke Functional Genomics Core, and members of the Horner Lab for valuable feedback and discussion. This work was supported by Burroughs Wellcome Fund (S.M.H) and National Institutes of Health grants R01AI155512 (S.M.H) and T32-CA00911 (H.S).

## Methods

### Cell culture

Huh7 cells, Huh7.5 cells, A549 cells, Vero cells, and 293T cells were grown in Dulbecco’s modification of Eagle’s medium (DMEM; Mediatech) supplemented with 10% fetal bovine serum (HyClone), 1X minimum essential medium non-essential amino acids (Thermo Fisher), and 25 mM HEPES (Thermo Fisher), referred to as complete DMEM (cDMEM). The identity of the Huh7 and Huh7.5 cells was verified by using the GenePrint STR kit (Duke DNA Analysis Facility). C6/36 cells were grown in Eagle’s minimum essential media (EMEM; ATCC) supplemented with 10% fetal bovine serum (HyClone), 25 mM N-2-hydroxyethylpiperazine-N′-2-ethanesulfonic acid (Thermo Fisher), and 1X nonessential amino acids (Thermo Fisher). Cells were obtained from the following sources: A549 cells, 293T, Vero cells, and C6/36 cells (CCL185, CRL-3216, CCL-81, and CCL-1660, respectively) from ATCC; Huh7 and Huh7.5 cells from Dr. Michael Gale Jr. (54). All cell lines were verified as mycoplasma free by the MycoStrip Mycoplasma Detection Kit (InvivoGen).

### Plasmids

The following plasmids were generated by insertion of PCR-amplified fragments from DENV Open Reading Frames (gift of Dr. Priya Shah) (41) into the KpnI-to-BstBI digested pEF-TAK-V5 using InFusion (Clontech): pEF-TAK-DENV-C-V5, pEF-TAK-DENV-pRM-V5, pEF-TAK-DENV-pRM-E-V5, pEF-TAK-DENV-NS1-V5, pEF-TAK-DENV-NS2A-V5, pEF-TAK-DENV-NS2B-V5, pEF-TAK-DENV-NS3-V5, pEF-TAK-DENV-NS4A-V5, pEF-TAK-DENV-NS4B-V5, and pEF-TAK-DENV-NS5-V5. The following plasmids were generated by insertion of PCR-amplified fragments into the XbaI-to-BamHI digested pLVX vector (Clontech): pLVX-Flag, pLVX-Flag-UFL1. The following plasmids were generated by insertion of PCR-amplified fragments into the EcoRI-to-BamHI digested pLVX vector: pLVX-Flag-UFM1 and pLVX-Flag-UFM1ΔC3.

### Antibodies

For immunoblotting, the following primary antibodies were used: R-anti-UFL1 (Novus Biologicals, NBP1-79039, 1:1,000), R-anti-UFBP1 (DDRGK1, Proteintech, 21445-1-AP, 1:1,000), R-anti-UBA5 (Abcam, ab177478, 1:1,000), R-anti-UFC1 (Abcam, ab189252, 1:1,000), R-anti-UFM1 (Abcam, ab109305, 1:1,000), M-anti-DENV NS3 (GeneTex, GT2811, 1:1,000), R-anti-ZIKV NS3 (GeneTex, GTX133320, 1:1,000), R-anti-DENV Capsid (GeneTex, GTX103343, 1:1,000), R-anti-ZIKV Capsid (GeneTex, GTX133317, 1:1,000), M-anti-Tubulin (Sigma-Aldrich, T5168, 1:1,000), Anti-V5-tag mAb-HRP-DirecT (MBL, M215-7, 1:5000), and M-anti-FlagM2-HRP (Sigma, A8592, 1:5000). For immunofluorescence microscopy, R-anti-Calnexin (Cell Signaling Technology, 2433S, 1:200), and M-anti-J2 (Cell Signaling Technology, 76651L, 1:200) were used.

### Cell line generation

UFM1 KO Huh7 cells were generated by using a purified ribonucleoprotein complex consisting of Cas9 protein and single-guide RNAs (sgRNAs) targeting UFM1 synthesized by Synthego. Cas9 protein and sgRNAs were mixed at a ratio of 1:6 and then added to 1×10^6^ Huh7 cells in Neon Resuspension Buffer R, followed by electroporation using the Neon Transfection System (Invitrogen). Following recovery, single cell clones were isolated and validated by anti-UFM1 immunoblot and genomic DNA sequencing, with one clone used here. Huh7-UFM1 KO cell pools overexpressing Flag-UFM1^WT^ or Flag-UFM1^ΔC3^ were generated by lentiviral transduction, as previously (55).

### Focus-forming assay for viral titer

Focus forming assays were performed similarly to previously described (56); briefly, supernatants were harvested from ZIKV or DENV-infected cells 48 h after infection, serially diluted, and used to infect naïve Vero cells in triplicate wells of a 48-well plate for 3 hours before overlay with methyl cellulose (Millipore Sigma, M0512). After 72 hours, cells were washed with phosphate buffered saline (PBS) and fixed with 1:1 methanol: acetone. Cells were blocked with 5% milk in phosphate buffered saline with 0.1% Tween (PBS-T), and then immunostained with M-anti-4G2 antibody generated from the D1-4G2-4-15 hybridoma cell line against the flavivirus envelope protein (ATCC; 1:2,000). Infected cells were visualized following incubation with a horseradish peroxidase–conjugated secondary antibody (1:500) and the VIP Peroxidase Substrate Kit (Vector Laboratories). The titer (focus-forming units (FFU) per milliliter) was calculated from the average number of 4G2-positive foci at 10X magnification, relative to the amount and dilution of virus used.

### Viral infections and generation of viral stocks

Infectious stocks of ZIKV-GLuc were generated by harvesting supernatant 3-5 days post-transfection of pCDNA6.2 MR766 single intron NS1 GLuc flanking HDVr (32) into 293T cells. The viral stocks were titered on Vero cells as described above. ZIKV^Flag-NS2A^ virus (gift of Dr. Matthew J. Evans) was generated similarly to previously tagged NS2A viruses (42), with a 3XFlag cloned into the junction between NS1 and NS2A in the plasmid-based rescue system for ZIKV MR766 (43). Infectious stocks of a cell culture-adapted strain of genotype 2A JFH1 HCV (57) were generated and titered on Huh-7.5 cells by focus-forming assay (FFA), as described. DENV (Dengue virus 2 Thailand/NGS-C/1944) (58), ZIKV (Zika virus/Homo sapiens/PRI/PRVABC59/2015), WNV (West Nile virus strain 3000.0259 isolated in New York in 2000) (59) and YFV-17D (Yellow fever virus 17D vaccine strain; gift of Dr. Helen Lazear) stocks were prepared in C6/36 cells and titered on Vero cells, as described above.

For viral infections, cells were incubated in a low volume of DMEM containing virus for 3-4 h, following which the infection media was replaced with cDMEM. The translation inhibitor cycloheximide (Sigma Aldrich, 100 µM) and the flavivirus RNA-dependent RNA polymerase inhibitor MK-0608 (Aldrich, 50 µM) were added to cells during infection and were included in the replacement media when indicated.

### Quantification of percent of virally infected cells

Cells were immunostained for either orthoflavivirual Envelope (M-anti-4G2, 1:1,000) or HCV NS5A (1:500; gift of Dr. Charles Rice), as well as nuclei (Hoescht). Percent of infected cells was calculated as the number of viral antigen positive cells / the number of total cells (4G2 or NS5A / DAPI) per field following imaging using a Cellomics ArrayScan VTI High Content Screening Reader (Duke Functional Genomics Facility). Values represent the mean ± SEM (n=4 fields) from three independent experiments, with >5,000 cells counted per experiment.

### *In vitro* transcription and electroporation of RNA

Plasmid DNA encoding a DENV replicon luciferase reporter (DENV-RLuc-SGR (8)), was linearized using XbaI (New England Biolabs). Purified linearized DNA was used as a template for *in vitro* transcription with the MEGAscript T7 transcription kit (Invitrogen). RNA was purified to be free of DNA and transfected in Huh7 cells via electroporation, as follows: 5 µg of RNA was mixed with 4×10^6^ Huh7 cells in Cytomix buffer (2 mM ATP, 10 mM K2HPO4, 0.15 mM CaCl2, 25 mM HEPES, 2 mM EGTA, 5 mM MgCL2, 120 mM KCl, 5 mM Glutathione) and electroporated at 27 V and 975 µF with a Gene Pulser XCell System (Bio-Rad). At 4 hours post-electroporation, cells were washed with PBS and cDMEM was replaced.

### Transfection

DNA transfections were performed using FuGENE6 (Promega) or PEIpro transfection reagent (Polyplus). The following siRNAs were used in this study: UFL1 (Qiagen-SI04371318), UFBP1 (Thermo Fisher-s35323), UBA5 (Qiagen-SI04146989), UFC1 (Qiagen-SI00755230), UFM1 (Horizon-L-021005-00-0005) or nontargeting AllStars negative control siRNA (Qiagen-1027280). siRNA transfection (30 pmol of siRNA; final concentration of 0.015 µM) was done using Lipofectamine RNAiMax (Invitrogen), with media changed 4 hours after transfection.

### Luciferase assay

Luciferase activity of GLuc or RLuc was measured using the *Renilla* Luciferase Assay System (Promega, E2810). Briefly, cell supernatant or cell lysate collected was collected in 1X *Renilla* luciferase lysis buffer. *Renilla* luciferase assay reagent was prepared by adding 1 volume of 100X *Renilla* luciferase substrate to 100 volumes of *Renilla* luciferase assay buffer. 20-50 µL of cell lysate or supernatant was plated into an opaque 96-well plate, and 100 µL of *Renilla* luciferase assay substrate was dispensed into each well, luciferase was read using a BioTek Synergy2 microplate reader.

### Immunoblotting

Cells were lysed in a modified radioimmunoprecipitation assay buffer (TX-100-RIPA) buffer (50 mM Tris [pH 7.5], 150 mM NaCl, 5 mM EDTA, 0.1% SDS, 0.5% sodium deoxycholate, and 1% Triton X-100) or NP40 lysis buffer (20 mM Tris [pH 7.4], 100 mM NaCl, 0.5% Nonidet P-40) supplemented with protease inhibitor (Sigma Aldrich) and Halt phosphatase inhibitor (Thermo Fisher) at 1:100, and post-nuclear lysates were isolated by centrifugation. Quantified protein, as determined by Bradford assay (Bio-Rad), was resolved by SDS/polyacrylamide gel electrophoresis (PAGE), transferred to nitrocellulose or polyvinylidene difluoride membrane membranes in the Trans-Blot Turbo buffer (Bio-Rad) using the Turbo-transfer system (Bio-Rad). Membranes were stained with Revert total protein stain (Licor Biosciences) and blocked with 3% bovine serum albumin (BSA) in PBS-T. Membranes were probed with primary antibodies directed against proteins of interest, washed with PBS-T, incubated with species specific horseradish peroxidase (HRP)-conjugate antibodies (Jackson ImmunoResearch, 1:5,000), or fluorescent secondaries (Licor Biosciences, 1:5,000), washed again with PBS-T, and treated with Clarity Western ECL substrate (Bio-Rad). Imaging was then performed using a LICOR Odyssey FC.

### Protein immunoprecipitation

Cells were lysed as above, quantified protein (between 100 and 500 µg) was incubated with anti-V5 magnetic beads (Cell Signaling Technology) or anti-Flag magnetic beads (Sigma Aldrich) in lysis buffer at room temperature for 45 minutes to 1 hour with head-over-tail rotation. The beads were then washed 3X in PBS or PBS-T and eluted in 2X Laemmli buffer (Bio-Rad) with 5% 2-Mercaptoethanol by incubating at 95°C for 5 minutes. Proteins were resolved by SDS/PAGE and immunoblotting as above.

### Immunofluorescence microscopy

Cells were fixed and permeabilized in 100% methanol and blocked with 10% FBS in PBS. Slides were stained with the indicated primary antibodies, washed 3X in PBS, incubated with conjugated Alexa Fluor secondary antibodies (Life Technologies), and mounted with ProLong Diamond + 4’, 6-diamidino-2-phenylindole (Invitrogen). Imaging was performed on a Leica DM4B widefield fluorescent microscope using a 63X oil objective. All images were processed with NIH Fiji/ImageJ.

